# High Level of Interaction between Phages and Bacteria in an Artisanal Raw Milk Cheese Microbial Community

**DOI:** 10.1101/2021.08.03.454940

**Authors:** Luciano Lopes Queiroz, Gustavo Augusto Lacorte, William Ricardo Isidorio, Mariza Landgraf, Bernadette Dora Gombossy de Melo Franco, Uelinton Manoel Pinto, Christian Hoffmann

## Abstract

Endogenous starter cultures are used in the production of several cheeses around the world, such as Parmigiano-Reggiano, in Italy, Époisses, in France, and Canastra, in Brazil. These microbial communities are responsible for many of the intrinsic characteristics of each of these cheeses. Bacteriophages are ubiquitous around the world, well known to be involved in the modulation of complex microbiological processes. However, little is known about phage–bacteria growth dynamics in cheese production systems, where phages are normally treated as problems, as the viral infections can negatively affect or even eliminate the starter culture during production. Furthermore, a recent metagenomic based meta-analysis has reported that cheeses contain a high abundance of phage-associated sequences. Here, we analyse the viral and bacterial metagenomes of Canastra cheese, a tradition artisanal cheese produced using an endogenous starter culture. We observe a very high phage diversity level, mostly composed of novel sequences. We detect several metagenomic assembled bacterial genomes at strain level resolution, and several putative phage-bacteria interactions, evidenced by the recovered viral and bacterial genomic signatures. We postulate that at least one bacterial strain detected could be endogenous to the Canastra region, in Brazil, and that its growth seems to be modulated by native phages present in this artisanal production system. This relationship is likely to influence the fermentation dynamics and ultimately the sensorial profile of these cheeses, with implication for all cheeses that employ similar production processes around the world.

## Introduction

Viruses are abundant in all ecosystems on Earth, presenting high genetic and taxonomic diversities, shaping biogeochemical cycles and ecosystem dynamics^1,2^. These obligate intracellular parasites are called bacteriophages (or simply phages) when they infect bacteria. The diversity and composition of phage-bacterial communities are influenced by environmental (*e*.*g*. pH, temperature, salinity) and biological factors^3,4^. Biological factors can be divided in host traits (species abundance, organismal size, distribution, physiological status, and host range) and virus traits (infection type, virion size, burst size, and latent period)^5^.

Many raw milk cheeses around the world are produced using endogenous starter cultures^6^, a complex microbial community composed of yeasts, bacteria and phage, all of which interact to create the final food product. This procedure often uses the backsloping method, where residual fermented whey is collected during production to be reinoculated as a starter culture on the next day’s production batch, effectively creating a continuous microbial growth system^7^. Brazil produces a wide variety of artisanal cheeses, several of which use the backsloping method^8,9^. Canastra cheese is one such cheese, and its endogenous starter cultures have recently been characterized using molecular techniques. They contain a diverse, but stable microbial community^10^. The community stability in these endogenous starters is assumed to be maintained by a continuous diversification process: when one species is excluded, the genetic function is kept present in the environment by another closely related species^11^. Studies of Canastra cheese microbiome are usually focused on the bacterial and fungal components^12,13^; however, little is known about its virome composition and interactions with the bacterial hosts.

Most bacteriophages found in dairy fermentation environments belong to the Siphoviridae family, such as 936, P335 and c2 groups. These non-enveloped viruses possess icosahedral morphology and non-contractile tails, with their genome encoded in double-stranded DNA (dsDNA), and commonly infect bacteria from the *Lactococcus* genus. Other less abundant phage groups are also able to infect bacteria from the genera *Leuconostoc, Lactobacillus, Streptococcus, Bacillus, Staphylococcus* and *Listeria*^14,15^. Phage-bacteria interactions tend to be highly specific, due to phage infection strategies that depend on the host binding proteins and the antiphage defense systems in the host^15,16^. These defense systems are classified in adaptive immune systems, including several types of CRISPR-Cas systems, and innate immune systems, such as restriction-modification (RM), abortive infection (Abi)^17^ and, most recently described systems BREX for Bacteriophage Exclusion^18^ and DISARM for Defense Island System Associated with Restriction-Modification^19^.

Here, we present the first description of the virome composition in Brazilian artisanal Canastra cheese and the phage-bacterial interactions in this food system. We identified 1,234 viral operational taxonomic units (vOTU) and explored the interactions with bacteria across seven cheese producing properties using a combination of viral and microbial metagenomic sequencing. We characterized a putative novel species of *Streptococcus* phage 987 group, as well as its potential host, a metagenome assembled genome (MAG) classified as *Streptococcus salivarius*. Finally, the relationships between 15 complete and high-quality phage genomes and 16 MAGs obtained from starter cultures and cheeses were evaluated.

## Results

### VLP-based description of the bacteriophage community present in the Canastra Cheese endogenous starter culture

We assessed the bacteriophage community present in the endogenous starter cultures used by seven artisanal Canastra Cheese producers in Brazil, located in São Roque de Minas and Medeiros, Minas Gerais, Brazil. Viral-like particles (VLPs) were enriched from these starter cultures using a 100 kDa filter membrane and used for metagenome sequencing. The sequencing reads were assembled and rigorously curated to remove bacterial DNA contaminants, producing a final viral sequence catalog (Fig. S1). Our final catalog yielded 908 complete and partial viral genomes, with contig sizes ranging from 1 × 10^3^ bp to 1.7 × 10^5^ bp and the coverage between 1.41 - 21108x with genomes classified as: complete (5), high-quality (12), medium-quality (23), low-quality (584), and not-determined quality (284) (Table S1).

Family level taxonomic classification for the viral catalog was made using the Demovir pipeline. The order Caudovirales (99%) prevailed, with only a minor number of sequences classified as Algavirales (0.55%) and Imitervirales (0.33%) (Table S1). Contigs were classified at family level as *Siphoviridae* (43.1%), followed by *Myoviridae* (12.1%), *Podoviridae* (3.4%), *Phycodnaviridae* (0.55%), *Mimiviridae* (0.33%), and *Retroviridae* (0.11%). The unassigned contigs corresponded to 40.3% (Fig. 1a). Integrase or site-specific recombinase genes were detected in 67 viral contigs (Fig. 1b) and VirSorter identified 7.38% of this viral sequence catalog as temperate phages (Fig. S2).

**Fig. 1.**
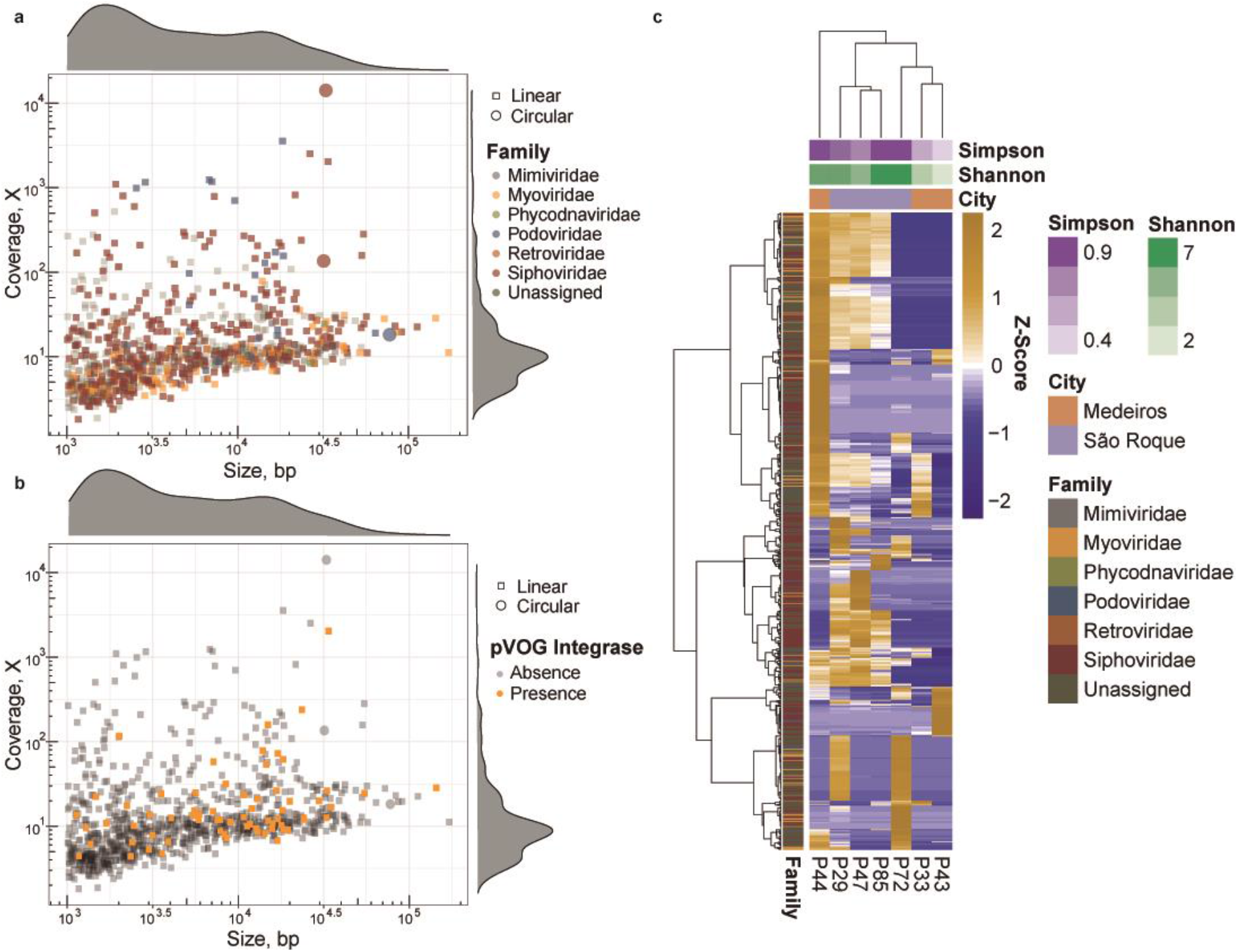
Viral diversity recovered by VLP metagenome sequencing. **a**, Size and coverage distribution of 908 putative viral genomes detected (coverage depth by genome length in bp), color coded by family classification. The families’ proportion were *Siphoviridae* (43.1%), *Myoviridae* (12.1%), *Podoviridae* (3.4%), *Phycodnaviridae* (0.55%), *Mimiviridae* (0.33%), *Retroviridae* (0.11%), and Unassigned (40.31%). **b**, Temperate phage distribution as detected by the presence of Integrase/Site-Specific Recombinase genes. **c**, Abundance distribution of viral genomes across starter samples analysed from 7 distinct cheese producers (viral contig abundances plotted as row z-score using normalized abundance values); only contigs present in at least 2 samples are shown in the heatmap (625 contigs).

Alpha diversity analysis of viral metagenomes was calculated using a normalized count table of reads mapped against the viral contigs. Four out of seven samples showed high values of diversity index (> 6 Shannon and > 0.94 Simpson), one sample showed medium values (4.66 Shannon and 0.78 Simpson), and two samples, low values (< 3 Shannon and < 0.6 Simpson) (Table S2). The sample with lowest diversity values (P43 sample, 1.29 Shannon and 0.39 Simpson) was dominated by one complete, high coverage viral genome (>21000x) belonging to the *Siphoviridae* family (Fig. 1c), and 94 contigs were classified at species level (Table S1; Supplementary Results)

### Classification and genome characterization of bacteriophages present in Canastra cheese endogenous starter culture

We used the vConTACT v2.0 clustering pipeline to refine the taxonomic assignment of our viral genome catalog^20^, using all known viral genomes. Only 2.75% of our contigs had any match against all viral genomes present in RefSeq, indicating a large amount of novel viral diversity (Fig. S3). Viral contigs clustered with RefSeq genomes belong to *Siphoviridae, Myoviridae* and *Podoviridae* families (Fig. 2a). Some complete and high-quality genomes were clustered with viral genomes present in RefSeq, such as phage ph.1.31871 and ph.1.31871 (VC148) and *Streptococcus virus* 9871, 9872, and 9874 that belong to phage 987 group. A phylogenomic analysis of this VC containing 98 *Streptococcus phages* genomes placed our novel genomes firmly within the 987 group. Considering that both phages presented < 95% ANI values compared to the four available genomes, we propose that phage ph.1.31871 and ph.1.31871 belong to a novel phage species within this group (Fig. S4). Although we observed a high similarity (as measured by ANI) between phages ph.1.32817 and ph.1.31872, they did not form a single vOTU, presenting an alignment fraction less than 85%, further indicating a potential strain differentiation.

**Fig. 2.**
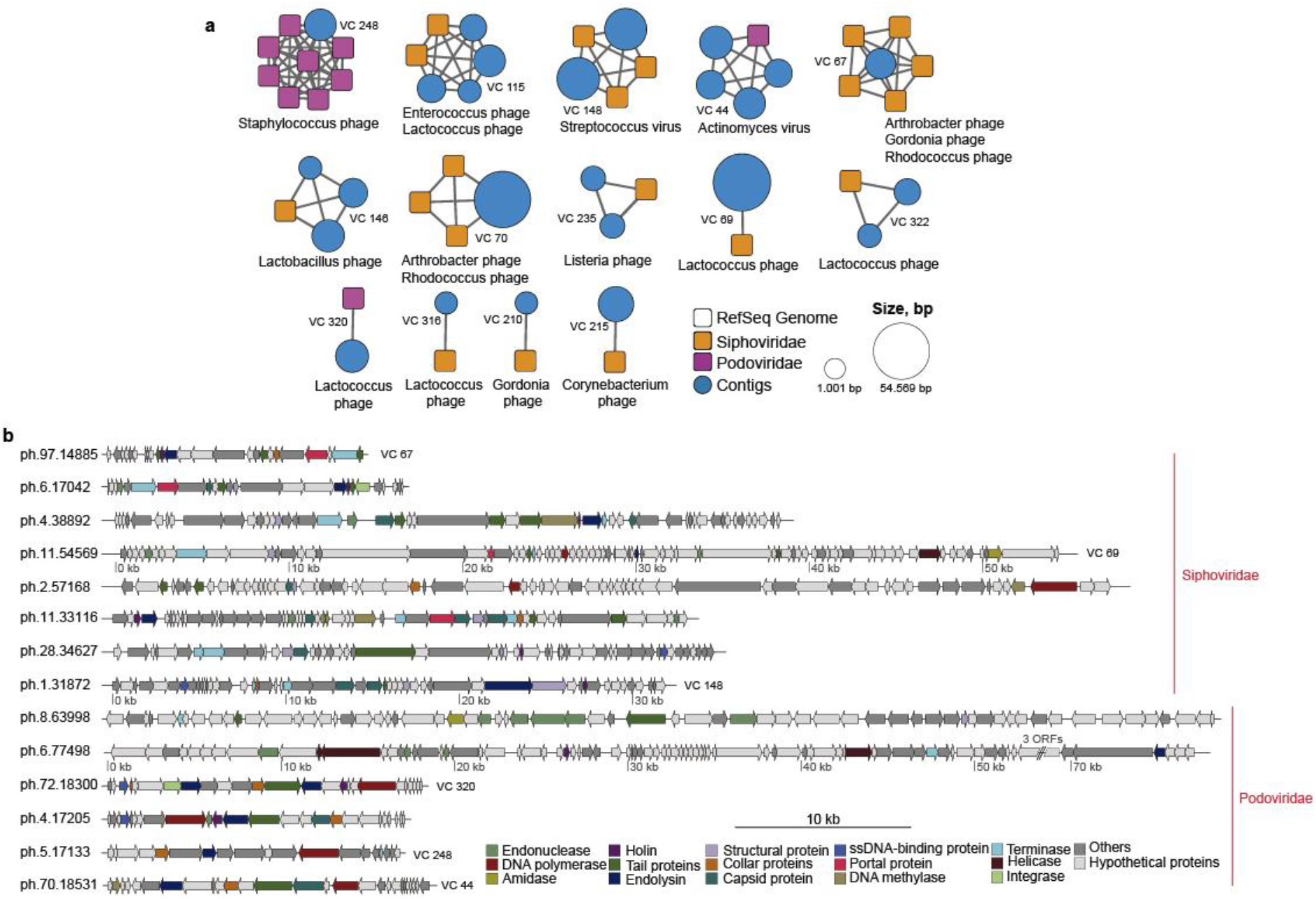
Viral Cluster taxonomy and genome annotation. **a**, Network of clusters formed between RefSeq genomes and viral contigs. RefSeq genomes and their family level classification are represented by squares and viral contigs by circles. Square colors indicate viral families and the size of circles represent the viral contig length in base pairs. **b**, Genome annotation of complete and high-quality bacteriophages genomes using pVOG database; some of these genomes were clustered with RefSeq genomes as shown by their VC number. Colors indicate the gene identification.

We annotate 14 of our 17 complete and high-quality putative viral genomes using the multiPhATE2 pipeline^21^ and the pVOG database (Fig. 2b). Eight genomes were classified as *Siphoviridae* with genomes sizes ranging from 14,885 to 57,168 bp, and six as *Podoviridae* with genomes sizes ranging from 17,133 to 77,498 bp. The most frequently annotated genes in these 14 viral genomes were Tail protein (22), Terminase (17), Endonucleases (15), Capsid protein (14), and Endolysin (13) (complete annotation is available in Table S3). We observed integrase genes in two genomes, including a fully recovered genome which clustered with *Lactococcus phage* asccphi28 belonging to P034 phage species (VC 320).

### Identification of lactic acid bacteria (LAB) in the endogenous starter cultures and cheese samples accessed by microbial metagenome sequencing

To better understand the microbial community and the interactions between phage and bacteria in Canastra cheese, we also carried out microbial metagenome sequencing using samples of endogenous starters and cheese produced with these same starters at 22-days of ripening, which is the minimal ripening time required by law in Brazil for the commercialization of Canastra cheese^22^. This data was initially analysed using MetaPhlAn2^23^, and the most abundant bacterial species detected were *Lactococcus lactis* (average of 30.6%), *Streptococcus thermophilus* (17.4%), *Streptococcus infantarius* (13.7%), *Streptococcus salivarius* (7.3%), and *Corynebacterium variabile* (5.5%) (Fig. S5). MetaPhlAn2 also detected *Lactococcus* phage ul36 (6.1%) in starter and cheese samples from producer P29 and in the starter from producer P72.

Having established this first characterization of the microbial community, we refined the bacterial species analysis by detecting metagenome-assembled genomes (MAGs). Metagenomic contigs were submitted to the Metagenomic Workflow of Anvi’o 6.1^24^. Contigs were binned using CONCOCT^25^ followed by manual curation. From a total of 50 MAGs obtained, we selected 16 refined MAGs with at least >50% completeness and <10% redundancy, nine of which showed more than 90% completeness (Table S4). FastANI^26^, CheckM^27^ and PhyloPhlAn 3.0^28^ were used to taxonomically classify each of these bins.

We identified 13 MAGs belonging to the Firmicutes phylum, two belonging to Actinobacteria, and one to Proteobacteria. The most representative genera were *Leuconostoc* (4 MAGs), *Streptococcus* (3 MAGs), and *Lactobacillus* and *Weissella*, (2 MAGs each), all of which had at least 95% of average nucleotide identity (ANI) to reference genomes. Other MAGs were classified as *Lactococcus, Rothia, Staphylococcus, Corynebacterium*, and *Escherichia* (Fig. 3a; Table S4). Additionally, MAG7 was assigned as *Streptococcus salivarius* and showed 94.2% of ANI to *Streptococcus salivarius* BIOML-A24 (genbank accession: GCF_009717045.1). A pangenomic analysis carried out with 95 *Streptococcus* genus reference genomes, and the phylogenomic tree constructed with 302 single-copy core genes, placed *S. salivarius* MAG7 between *S. salivarius* and *S. thermophilus* groups, further indicating its potential as a new strain (Fig. S6).

**Fig. 3.**
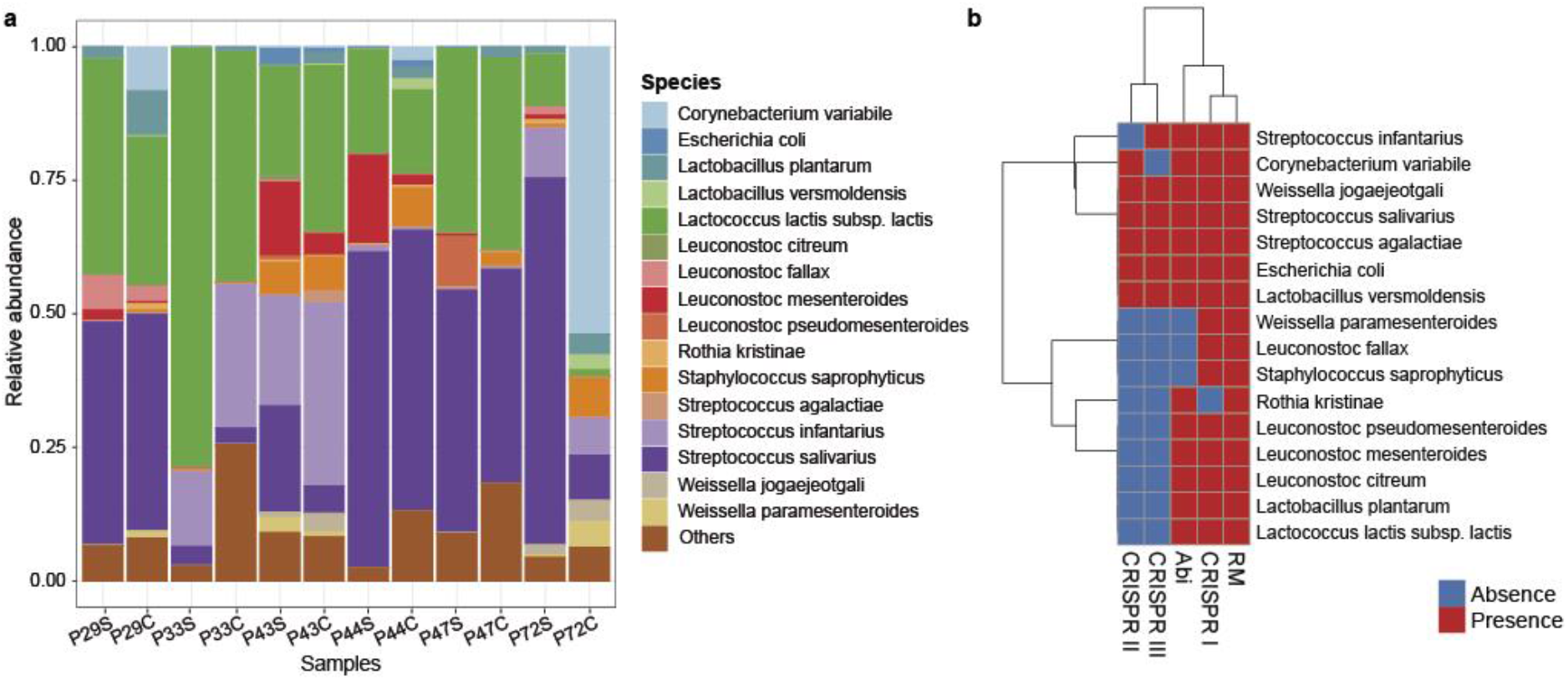
Metagenome-assembled genomes (MAGs) composition and antiphage defense mechanisms in endogenous starter and cheese. **a**, Relative abundance of each MAG classified at species level in 12 paired starter and cheese samples. Others represent reads mapped against lower quality MAGs. **b**, Presence and absence of antiphage defense mechanisms found in each MAG.

The most abundant MAGs across all samples were classified as *Streptococcus salivarius, Lactococcus lactis* subsp. *lactis, and Streptococcus infantarius* (averages of 34.6%, 33%, and 11.4% respectively), detected in all samples. An inverse relationship was detected between *S. infantarius* e *S. salivarius* abundances across all studied samples. We observed a high level of similarity between the community composition observed in the starter and cheese samples within the same producer (Wilcoxon test p value = 0.001, comparison of within versus between producers Bray-Curtis distances), with some species decreasing or increasing in relative abundance, such as the decrease of *L. lactis* subsp. *lactis* and increase of S*S. infantarius* in samples from producer P33. A departure from this tight within-producer relationship between starter and cheese samples was observed only for producer P72, where we observe a marked increase in *Corynebacterium variabile* relative abundance from starter (0.003%) to cheese (56.9%).

### Antiphage defense mechanisms found in Canastra cheese MAGs

The 16 MAGs were screened for known bacterial defense system genes and manually filtered for specific antiphage defense mechanisms. We identified a total of 395 defense genes belonging to restriction modification (RM), abortive infection (Abi), CRISPR-type I, II and III mechanisms (Fig. 3b). The *Rothia kristinae* genome had no CRISPR-cas system genes and all *Leuconostoc* genomes presented only the CRISPR-I system. *Weissella jogaejeotgali, Streptococcus salivarius, Streptococcus agalactiae, Escherichia coli*, and *Lactobacillus versmoldensis* harbored all five types of defense genes. Another five MAGs harbored three types of defense genes, such as *Lactococcus lactis* subsp. *lactis* and *Leuconostoc mesenteroides*, and four MAGs had only two types of defense genes, *Weissella paramesenteroides, Leuconostoc fallax*, and *Staphylococcus saprophyticus* harbored RM and CRISPR-type I, and *Rothia kristinae* with RM and Abi systems. We did not find any defense genes classified as DISARM, BREX, Thoeris, Shedu, Gabija, and others (Table S5).

### Presence of CRISPR spacers and prophages in MAGs

The phage infection history of a given bacterium can be studied by characterizing CRISPR arrays present in their genome. Here, we identified CRISPR spacers in our MAGs using PILAR-CR^29^, and each spacer consensus generated was matched against our viral contig catalog and the IMG/VR database. A total of 10 CRISPR arrays were found in 4 of the 16 MAGs (Supplementary Data). The MAG classified as *Streptococcus salivarius* (MAG7) harbored five arrays with 153 spacers and average length of 212 bp. The second MAG with most CRISPR arrays was MAG3, classified as *Escherichia coli*, which harbored three arrays with 106 spacers and 267 bp of average length. The other two MAGs (*Lactobacillus versmoldensis* and *Weissella jogaejeotgali*) harbored only one array each.

All CRISPR arrays of *Streptococcus salivarius* matched (more than 95% identity) with phages from IMG/VR database, represented by phages from families *Siphoviridae* and *Myoviridae*, and having genera *Streptococcus* and *Streptococcus thermophilus* as predicted host lineages. The array present in the *Weissella jogaejeotgali* MAG matched with phages from family *Siphoviridae*, and no match was found for the array present in the *Lactobacillus versmoldensis* MAG (Table S6). Two of three arrays detected in the *E. coli* MAG presented sequence signatures for *Siphoviridae, Myoviridae, Podoviridae*, and *Inoviridae* (Tubulavirales) phage families, all of which have *Escherichia, Klebsiella*, and *Xanthomonas* species as predicted host lineages. Only one array, detected in the *E. coli* MAG, matched with phage contigs from our viral catalog, one of these was the ph.11.54569 classified as *Lactococcus* phage and grouped within VC69 (Fig. 2). We also identified four intact prophage sequences in our MAGs (Table S7; Supplementary Results).

### Phages contigs recovery from microbial metagenomes

It was possible to recover phage sequences not detected in the VLP-isolated dataset, by analysing the microbial metagenome dataset using VirSorter^30^. A total of 514 viral contigs were identified from the 12 metagenome samples (Table S2), and their taxonomic classification at family level revealed a prevalence of *Siphoviridae* (85%), followed by *Myoviridae* (4,2%), and *Podoviridae* (2,5%). We recovered 2 complete and 10 high-quality genomes, while the remaining genome fragments were of medium-quality (27), low-quality (467), and not-determined (8). The two complete phage genomes recovered from the bacterial metagenomes were the same as found when sequencing VLPs obtained from producers P33 and P43 endogenous starter samples (ph.1.31872 and ph.1.32817).

### Correlations between phage and MAG populations in cheese metagenomes

We expanded our viral catalog to include the new phage detected using the metagenome dataset, by comparing all contigs obtained from the viral and bacterial metagenomes using FastANI, creating a final virus list with 1234 unique phage contigs. Then, we mapped all reads obtained from each sample to this expanded viral list to create a viral OTU (vOTU) table for further comparative analysis. A Procrustes analysis of Jensen-Shannon divergence matrices did not indicate a correlation between viral and bacterial communities in starter samples (0.702 with p = 0.39, Fig. 4a); however, the correlation between the viral and bacterial communities in the cheese samples was significant (0.865 with p = 0.0005, Fig. 4b). We also identified a significant correlation between bacterial communities present in starter versus cheese samples, but not for phage communities (Fig. S7).

**Fig. 4.**
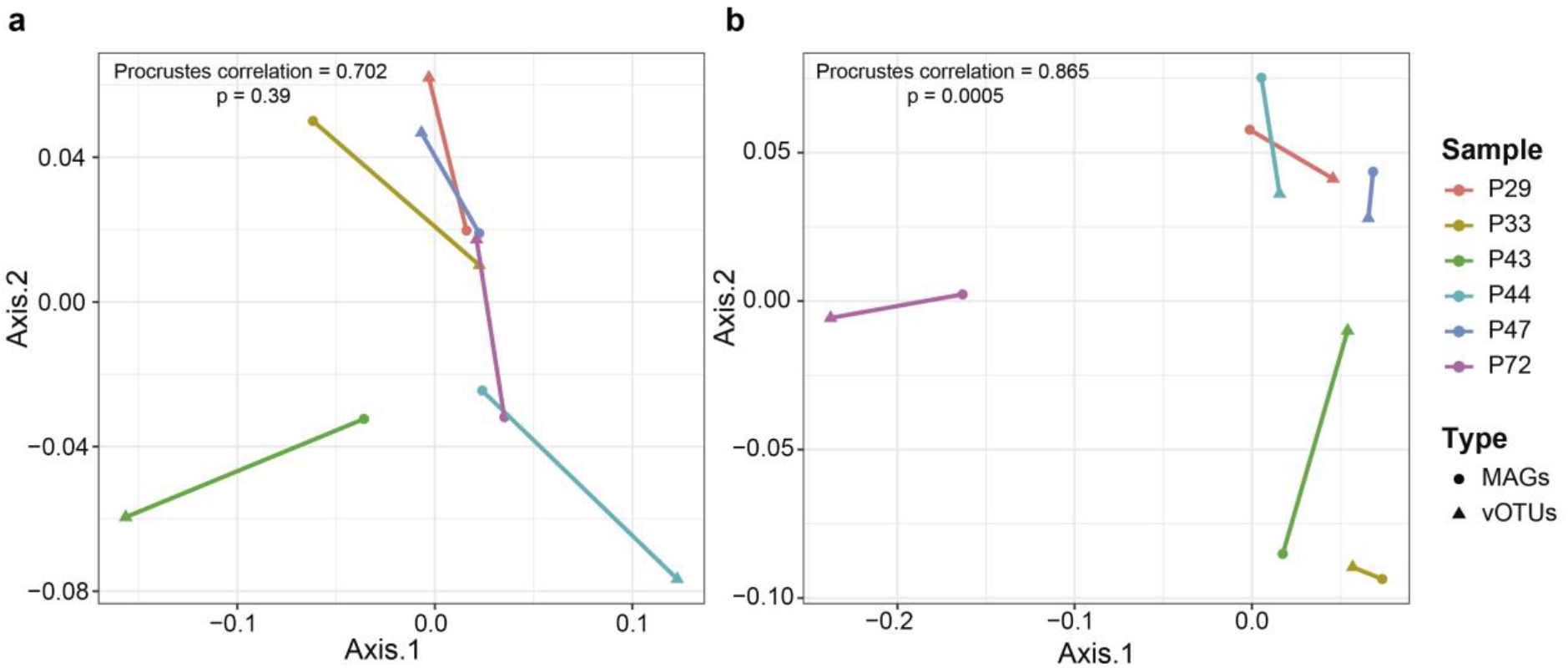
Procrustes analysis using PCoA coordenations of viral and bacterial communities. **a**, Correlations between viral (vOTUs = 1234) and bacterial communities (MAGs = 16) in endogenous starter cultures samples. **b**, Correlations between viral and bacterial communities in cheese samples. Distance matrices were calculated using Jensen-Shannon divergence in both cases.

### Evidence of interactions between phages and bacteria during cheese production

We further evaluated the existing relationship between the 15 high quality phage genomes and 16 high quality bacteria MAG in the Canastra cheese production system, using Spearman correlations based on normalized phage/bacteria abundances and on predicted host infection using the software WIsH^31^. We considered that phages present in the initial starter culture would infect their hosts during cheese production, and therefore a negative correlation was expected. Indeed, negative correlations were found both in endogenous starters and in cheese samples (Fig. 5 and Table S8). For instance, phages ph.1.32817 (ρ = −0.84, *p* = 0.03) and ph.1.31872 (ρ = −0.88, *p* = 0.02) both showed negative correlations with *S. salivarius* (MAG7), and phage ph.1.31872 was also negatively correlated with *L. pseudomesenteroides* (MAG8) (ρ = −0.83, *p* = 0.04). We also observed negative correlations between phages ph.11.33116 and *L. lactis* subsp. *lactis* (MAG4) (ρ = −0.84, *p* = 0.03); phages ph.8.63998 and *L. mesenteroides* (MAG10) (ρ = −0.88, *p* = 0.01); and ph.2.57168 and *S. agalactiae* (MAG14) (ρ = −0.88, *p* = 0.01). Additionally, the predicted host for phages ph.8.63998 and ph.2.57168 was *L. lactis* subsp. *lactis* (MAG4) (null model *p-value* < 0.05), and the two phages and MAG4 were present in all samples. Other predicted hosts were *S. saprophyticus* (MAG1), for phage ph.5.17133, and *C. variabile* (MAG5) for phages ph.6.17042 and ph.97.14885 (Table S9). Although an overall correlation pattern could be observed across all samples, we also detected a high level of individual variation across all analysed producers, indicating a high level of producer specialization (Fig. 5 and Fig. S8).

**Fig. 5.**
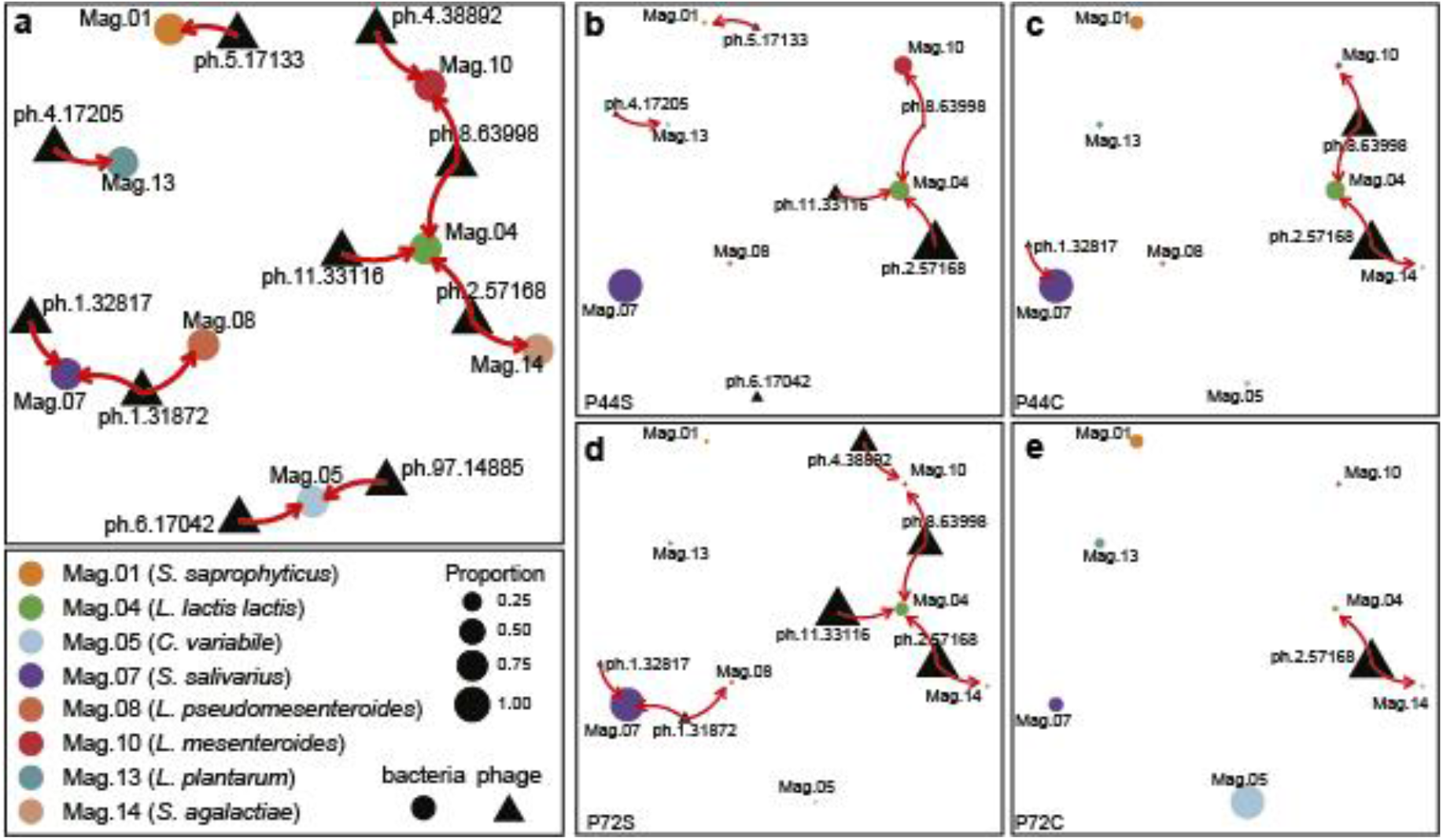
Phage-bacteria interaction network during cheese production. **a**, General network describing the putative phage-bacteria infection interactions present across all samples. **b-e**, Examples of putative phage-bacteria infection interactions for two distinct producers (P44: b-c; and P72: d-e), and for separate starter and cheese samples (starter: b and d; cheese: c and e). Nodes represent each MAG and phage analysed, edges represent putative infection relationships between phage and bacteria.

## Discussion

Endogenous starter cultures are used in the production of several cheeses around the world, such as Parmigiano-Reggiano, in Italy, Époisses, in France, and Canastra, in Brazil. The bacterial composition of these starters is often well characterized, but little is known about their phage–bacteria growth dynamics, with phages normally treated as problems, as viral infections can negatively affect or even eliminate the starter culture during production. Only a few studies characterize the bacteriophage community composition in cheese samples using viral metagenomes, mostly using whey or cheese rind samples^32,33^, and a recent metaanalysis using 184 cheese microbial metagenomes identified a high abundance of phage-associated sequences^34^. Here, we report the first study of phage-bacteria dynamics present in an artisanal cheese produced using the backslopping method, as well as its endogenous starter culture, from seven distinct artisanal Canastra Cheese producers in Minas Gerais Gerais state, in Brazil.

We sequenced viral and microbial metagenomes, recovering a total of 1,234 vOTUs, including 18 high-quality or complete viral genomes, and 16 metagenome assembled bacterial genomes (MAGs). The majority of the viral genomes were assigned to *Siphoviridae, Myoviridae*, and *Podoviridae* families, which are commonly encountered in dairy systems^14,15,32^. We have also observed a high proportion of unclassified contigs or classified only at family level, even when they were considered complete and high-quality contigs. There is a high level of viral diversity variation across all analysed starter samples, with differences as high as 5-fold being observed for Shannon diversity index. Samples with low viral diversity tended to be dominated by two phages that clustered with *Streptococcus* virus group 987 reference genomes. This phage group has been recently discovered and described as a novel emerging group of *S. thermophilus* phages. Group 987 phage is thought to have originated from genetic exchange events between *L. lactis* P335 phage group and *S. thermophilus* phages, acquiring morphogenesis related genes and replication modules from each group respectively^35^. We are presenting the first description of phage 987 group in a Brazilian cheese, and the observation of a putative novel phage species in this group.

One of our most unique findings is the detection of a complete genome, phage ph.72.18300, which clustered with *Lactococcus* phage asccphi28 genome, belonging to the group P034 phage species (family *Podoviridae*), a group rarely found in the dairy industry^14,36^. *Lactococcus* phage asccphi28 shows more genetic and functional similarities with phages normally infecting *Bacillus subtilis* and *Streptococcus pneumoniae* than other *Lactococcus lactis* phages^36^. Our analysis indicates that this phage is potentially a novel viral species within the *Lactococcus* phage asccphi28 group. Another phage genome, ph.5.17133, was classified as a *Staphylococcus* phage belonging to the genus *Rosenblumvirus* (*Podoviridae*), a phage group currently being considered for its potential use in bacteriophage therapy in veterinary medicine, as a means to treat *Staphylococcus*-positive mastitis^37,38^.

Recent studies have highlighted the importance of characterizing microbial strains within dairy-related systems to understand microbiome assembly and function in several habitats, such as cheese rinds^34,39,40^. For instance, distinct bacterial strains can respond to environmental and biological stress in different ways. We have characterized 16 MAGs at strain level resolution, including LAB such as *Lactococcus lactis* subsp. *lactis, Streptococcus salivarius*, and *Streptococcus infantarius*, followed by less abundant strains of *Leuconostoc, Lactobacillus*, and *Weissella*. We also found a large amount of evidence for the interaction between phage and bacterial strains in Canastra cheese production system, with all MAGs showing at least two types of antiphage defense systems, such as CRISPR, restriction-modification (RM) and abortive infection (Abi). The presence of several methyltransferase genes was observed within the phage contigs, which is a well-known phage evasion mechanism against bacterial RM systems^41,42^. Furthermore, we also identified CRISPR spacer arrays in 4 of the 16 analysed MAGs, indicating previous infections and active evolution of the adaptive immune system in these strains. The MAGs identified as *S. salivarius* and *E. coli* showed multiple spacers from different phage species, suggesting multiple infection events^43^. Additionally, our results demonstrate that *Streptococcus salivarius* MAG7 is likely a new strain. Therefore, we postulate that the MAG 07 *Streptococcus salivarius* strain could be endogenous to the Canastra region, in Brazil. Furthermore, its growth seems to be modulated by native phages present in this artisanal production system, and this relationship is likely to influence the fermentation dynamics and ultimately the sensorial profile of these cheeses.

We observed a high level of similarity between proportions of predominant microbial species, from starter to cheese samples, indicating a resilient microbial ecosystem. This also highlights that although microorganisms could be acquired during the cheese production and ripening stages, the overall microbiome composition and structure present in Canastra cheese is primarily determined by the starter culture. Nevertheless, one producer seemed to depart from this pattern in our sampling, where *Corynebacterium variabile* dominated the cheese samples while being a minor component of the starter culture. *Corynebacterium variable* is found in smear-ripened cheeses and is responsible for flavor and textural properties during ripening process and strains of this species are known to compose the microbiome of surface cheeses^44^. It is possible that *C. variable* is acquired during the maturation process, when the cheese is in direct contact with several surfaces for a prolonged period of time.

Phages ph.1.32817 and ph.1.31872 are negatively correlated with *S. salivarius* MAG7, potentially indicating their ability to infect this species. *Streptococcus salivarus* MAG7 was the most abundant *Streptococcus* species in absence of 987 phage strains, and when these phages were present, *S. infantarius* MAG6 became the dominant *Streptococcus* species. Thus, the control of *S. salivarius* by the 987 phage provides an adaptive advantage to *S. infantarius*, allowing it to become the dominant *Streptococcus* species in this lactic fermentation ecosystem. Phages belonging to the 987 group are also described as being able to infect some *Lactococcus* species, however we did not observe any evidence for this interaction in our analysis.

For cheese samples, it was possible to detect a global relationship between the composition of phages and bacteria. Biochemical and environmental changes occurring during cheese production and ripening, such as pH and salinity, are known to affect microbiome competition in many types of cheese^6,8^. Thus, it is likely that these same factors could influence phage-bacterial interactions. Other factors intrinsic to lactic fermentation systems, and that can influence phage-bacteria interactions, are the metabolism of residual lactose, lactate, citrate, and lipolysis and proteolysis^45,46^. Previous studies have demonstrated that there is a balance between active starter cells and lysed cells to control the lactose degradation and proteolysis, respectively, and some milk proteins can interfere in phage activity^47,48^. The matrix structure of cheese can also impact those interactions by firmness and viscosity of cheese^48^. Finally, phage dispersion rates in a matrix of hard cheese will be different from that in soft cheese or in starter culture and whey solutions.

In conclusion, this study revealed a rich and diverse phage population permeating all cheese and endogenous starter culture samples, with high levels of phage-bacteria interactions in the Canastra cheese production system. We extensively described the main phage members of these microbial ecosystems, and identified a likely novel phage species, belonging to *Streptococcus* phage 987 group, as well as its putative host, a novel strain of *S. salivarius*. We observed a dynamic yet stable microbial ecosystem during cheese production, marked by genomic evidence of continued phage-bacteria interactions, where general patterns emerged, yet maintaining a high level of inter-producer variability. This is a first effort to describe and understand the viral composition and ecological dynamics within the Canastra Cheese production system. We provide a solid background for further mechanistic studies focused on identifying phage-bacteria interactions in artisanal cheeses, with likely impact in similar production systems around the world.

## Methods

### Sampling

Samples were collected from seven cheese producers located in São Roque de Minas and Medeiros cities, in the Serra da Canastra region, state of Minas Gerais, Brazil. For the viral metagenome analysis, 50 mL of the endogenous starter culture intended to be used in the daily production and originally obtained from the previous day of production (backsloping method), were sampled and aliquoted in sterile polypropylene tubes. Cheeses produced with these starters were also sampled, at 22-days of ripening, when the cheeses are initially released for sale and human consumption. All samples were placed at −20 °C for the duration of the field trips (as long as 48h), and shipped frozen overnight to the laboratory where they were stored at −80 °C until further processing.

### Virus-like particles (VLPs) concentration

VLPs were obtained by filtering and centrifugation. Briefly, 50 mL of starter culture was vortexed for 60s, and centrifuged at 2500G speed for 10 min. The supernatant was transferred to a new tube and the pH adjusted to ∼4.6 with 1M HCl or 1M NaOH as needed^32^. Adjusted samples were sequentially filtered using 0.45 and 0.22 µm Millex®–HV PVDF syringe filters (Merck Millipore, Tullagreen, Cork, IRL). Filtered samples were centrifuged at 5000G using an Amicon ^®^ Ultra 15 (100 kDa) (Merck Millipore, Tullagreen, Cork, IRL) following the manufacturer’s protocol (30 min of centrifugation for every 12 ml). The final concentrated solution was diluted on the Amicon Filter with SM buffer (NaCl 100 mM, MgSO4•7H2O 8 mM M, Tris-Cl 50 mM pH 7.5), and centrifuged at 5000G up until ∼ 3 ml of the concentrated phage solution was recovered (complete protocol are available in Supplementary Information).

### DNA Extraction

The concentrated VLP stocks were pH adjusted to 7.5 using HCl of NaOH, if needed, prior to DNA extraction. VLP DNA extraction was made following the protocol produced by Jakočiūnė and Moodley^49^ with the DNeasy Blood & Tissue Kit (Qiagen Inc.). Briefly, 450 µl of phages concentrated were incubated with 50 µL of DNase I 10x buffer, 1 µL DNase I (1 U/µL), and 1 µL RNase A (10 mg/mL) (Sigma-Aldrich, St. Louis, MO, USA) for 1.5 h at 37 °C. DNase and RNase were inactivated with 20 µl of EDTA 0,5M (Sigma-Aldrich, St. Louis, MO, USA) (final concentration of 10 mM) for 20 min at room temperature. To digest phage protein capsid, 1.25 µL of Proteinase K (20 mg/mL) (Invitrogen, Waltham, MA, EUA) was added and incubated for 1.5 h at 56 °C without agitation. DNA purification was carried out using DNeasy Blood & Tissue Kit with 500 µL of lysed phage to increase the yield of extracted DNA.

Total DNA was extracted using the E.Z.N.A.^®^ Soil DNA Kit (Omega Bio-tek, Inc., Norcross, GA, USA): 250 mg of rind cheese was used for the DNA extraction; 3 mL of starter culture was centrifuged, and the cell pellet was used for extraction according to manufacturer’s instructions. The integrity of the DNA extracted was evaluated by electrophorese. DNA was quantified using Quant-iT™ PicoGreen™ dsDNA Assay Kit (Thermo Scientific, Waltham, MA, USA) and Synergy H1 Hybrid Reader (BioTek, Winooski, VT, USA).

### Sequencing

The Nextera DNA Flex Library Prep Kit (Illumina, San Diego, CA, USA) was used to generate dual-indexed paired-end Illumina sequencing libraries following the manufacturer’s instructions. Libraries were sequenced using 2 x 150 nt paired-end sequencing runs (4 lanes on separate runs) on NextSeq Genome Sequencer (Illumina) with a NextSeq 500/550 High Output Kit v2.5 at Core Facility for Scientific Research – University of Sao Paulo (CEFAP-USP).

### Viral metagenome bioinformatics analyses

The quality of raw sequences was verified using FastQC v0.11.9^50^. NextSeq adapters were removed using BBDuk (BBTools, https://jgi.doe.gov/data-and-tools/bbtools/) with the following parameters: ktrim=r k=23 mink=11 hdist=1. The quality trimming was also done using BBDuk with parameters: qtrim=r trimq=10 minlen=60 ftr=139. To assemble the quality filter reads of each viral metagenome we used SPAdes 3.15.0^51^ with metagenomic function (metaspades.py) and automatic parameters of kmers sizes. Generated contigs were filtered to remove short and redundant sequences using BBMap function dedupe.sh with parameters: minscaf=1000 sort=length minidentity=90 minlengthpercent=90. Open reading frames (ORF) were predicted using Prodigal v2.6.3^52^ in metagenomic mode. The final catalog of viral contigs was generated using similar analysis and criterion of Shkoporov et al.^53^, with some adaptations. Briefly, the search for amino acid sequences of predicted proteins we used a Hidden Markov Model (HMM) algorithm (hmmscan from HMMER v3.3) against HMM database of prokaryotic viral orthologous groups (pVOG)^54^ considering the significant hit e-value threshold of 10^−5^. Ribosomal proteins were searched using Barnnap 0.9 (https://github.com/tseemann/barrnap) with an e-value threshold of 10^−6^. Contigs were aligned against the viral section of NCBI RefSeq database using BLASTn^55^ of BLAST+ package^56^ with following parameters: e-value < 10^−10^, covering > 90% of contig length and > 50% identity. We also used VirSorter v1.0.6^30^ as criteria to predict viral sequences with its standard built-in database of viral sequences, with parameter: --db 1. Contigs that meet at least one of the following criteria were included in the final catalog of viral sequences: 1) VirSorter Positive, 2) BLASTn alignments to viral section of NCBI RefSeq, 3) minimum of three ORFs producing HMM-hits to pVOG database, and 4) be circular. Contigs selected at filter step (n = 908) were taxonomic assignment using Demovir script (https://github.com/feargalr/Demovir) with default parameters and vConTACT v2.0^20^clustering pipeline, a network-based analytical tool that uses whole genome gene-sharing profiles and distance-based hierarchical clustering to group viral contigs into virus clusters (VCs). Besides our viral contigs, we also included in the pool known viral genomes (NCBI RefSeq database release 88). Integrase and site-specific recombinase genes were identified in HMM hit to pVOG annotation of viral contigs. A counting table of viral contigs was generated, mapping unassembled sequences from each library using BBMap with the following parameters: minid=0.99 ambiguous=random. Reads count to contigs with coverage values less than 1X for 75% of a contig length, were set to zero^57^. The number of sequences mapped on viral contigs was normalized using the DESeq2 package^58^. Completeness, contamination and quality of contigs were assessed using CheckV^59^. We selected 14 contigs classified as Complete and High-quality genomes, annotated them using the multiPhATE2 pipeline^21^, with ORF prediction by Prodigal, gene annotation with hmmscan using pVOG database.

A phylogenomic tree of *Streptococcus* phages (complete list of phage genomes and code accession are available in Table S10) was constructed, based on previously study of Philippe et al.^60^, using VICTOR^61^: Virus Classification and Tree Building Online Resource. Briefly, pairwise comparisons of the nucleotide sequences were realized using the Genome-BLAST Distance Phylogeny (GBDP) method^62^ under settings recommended for prokaryotic viruses. The intergenomic distances were used to infer a balanced minimum evolution tree with branch support via FASTME 2.0^63^ and branch support was inferred from 100 pseudo-bootstrap replicates each and visualized with FigTree 1.4.4.

### Microbial metagenome bioinformatics analyses

The quality of raw sequences was verified using FastQC v0.11.9. NextSeq adapters were removed using BBDuk (BBTools) with the following parameters: ktrim=r k=23 mink=11 hdist=1. The quality trimming was also done using BBDuk with parameters: qtrim=r trimq=10 minlen=100 ftr=140. Compositional profiles of microbial metagenomes samples were assessed using MetaPhlAn2 (v2.7.5)^23^. To assemble the quality filter reads of each microbial metagenome we used SPAdes 3.15.0^51^ with metagenomic function (metaspades.py) and automatic parameters of kmers sizes. Generated contigs were filtered to remove short and redundant sequences using BBMap function dedupe.sh with parameters: minscaf=1000 sort=length minidentity=90 minlengthpercent=90.

We examined strain level metagenome-assembled genomes (MAGs) by co-assembling quality filtered sequences using MEGAHIT assembler^64^, with parameter --presets meta-large, and the contigs generated were filtered using BBMap function dedupe.sh with parameters: minscaf=1000 sort=length minidentity=90 minlengthpercent=90. The resulting filtered contigs were submitted to Metagenomic Workflow using Anvi’ 6.1^24^. Briefly, we created contig databases, mapping samples reads against contigs using Bowtie2^65^ and converted SAM files to BAM with SAMtools^66^; sequence homologs were searched and added to contigs database with hidden Markov Model (HMM) using HMMER^67^; genes were annotated functionally using NCBI’s Clusters of Orthologus Groups^68^and taxonomically using Centrifuge^69^; we created an anvi’o profile database with contig length cutoff of 2,500 bp; contigs binning were made using CONCOCT software^25^ and generated bins refined manually using anvio-refine function. We selected 16 refined MAGs following the criterias of >50% of completeness and <10% of redundancy (MAGs contigs are available in Supplementary Data). Taxonomic inference of MAGs was done using CheckM^27^ and PhyloPhlAn 3.0^28^ with database SGB.Nov19, after that, all complete sequences of each MAG species from NCBI RefSeq were downloaded and compared them with MAG sequences using FastANI^26^. A counting table of viral contigs was generated, mapping unassembled sequences from each library using BBMap with following parameters: minid=0.99 ambiguous=random. The number of sequences mapped on MAGs contigs was normalized using the DESeq2 package. Pangenomic analysis of *Streptococcus* genus was performed with 94 complete genomes (Table S11) selected from previous study by Gao et al.^70^ and MAG7 using Anvi’o v6.1 with the pangenomic workflow. Briefly, an anvi’o genome database was created, computing (with flag: --use-ncbi-blast; and parameters: --minbit 0.5 -- mcl-inflation 8) and displaying pangenome.

CRISPR spacers of MAGs were identified using PILAR-CR^29^, the spacers consensus generated were matched with our viral contig catalog and the IMG/VR database using BLASTn of BLAST+ package with the following parameters: -qcov_hsp_perc 80 -task blastn -dust no -soft_masking_false^71^. Matches of > 90% sequence identity for viral contig catalog and > 95% identity for IMG/VR database were considered. The antiphage defense mechanisms of MAGs were detected following the methods shown by Bezuidt et al.^71^. Briefly, we screened predicted genes for domain similarity of known defense systems against the conserved domains database (CDD) of clusters of orthologous groups (COGs) and protein families (Pfams) using RPS-BLAST (e-value < 10^−2^)^55^. The results were manually filtered for the identification of phage-specific defense systems (complete list is available in Supplementary Methods). Prophages present in MAGs were detected using the PHASTER web server^72^.

Phage contigs present in microbial metagenomes were assessed using VirSorter v1.0.6^30^ as criteria to predict viral sequences with its standard built-in database of viral sequences, with parameter: --db 1. We selected contigs classified only in categories 1, 2 (Phages), 4 and 5 (Prophages) of VirSorter output (n = 514). Open reading frames (ORF) were predicted using Prodigal in metagenomic mode. Those contigs were submitted to the same process that viral metagenome contigs, ORFs annotation with pVOG, ribosomal proteins searched using Barnnap and contigs aligned against the viral section of NCBI RefSeq database using BLASTn. In this case, we used VirSorter Positive as the only criterion to include in the final catalog of viral contigs from microbial metagenome sequences. The taxonomic assignment of contigs, completeness, contamination, and quality of contigs were also made using Demovir, vConTACT v2.0 and CheckV with the same parameters used to viral metagenome sequences.

Our vOTU table of contigs was created combining the two viral contigs catalogs (viral and microbial metagenome). We compare the Average Nucleotide Identity (ANI) of contigs from viral and bacterial metagenomes within and between them using FastANI, with criterion of ANI≥ 95% and minFraction>85%^73^. A counting table of vOTU contigs was generated mapping unassembled sequences from each library using BBMap with following parameters: minid=0.99 ambiguous=random. Read counts to contigs with coverage values less than 1 x for 75% of a contig length, were set to zero^57^. The number of sequences mapped on viral contigs was normalized using the DESeq2 package.

## Statistical analysis

All analyses were carried out using the statistical software R^74^ and specific packages as follows: we estimated alpha-diversity using Shannon (log base 2) and Simpson diversity indexes, and richness using the number of observed viral OTU’s for each sample using packages *vegan*^75^ and *microbiome*^76^ packages. We compared similarities between samples of viral and microbial metagenomes through Principal Coordinates Analysis (PCoA) using Jensen-Shannon divergence, followed by Procrustes analysis using *phyloseq*^77^ package. Correlations between normalized abundance values of vOTUs present in starter culture versus normalized MAG abundances present in the cheese were calculated to explore potential phage-bacteria predation relationships. Correlations were deemed significant if they had a value equal to or lower than −0.8 and p value <= 0.05. Additional phage bacterial relationships were explored using WiSH^31^ with a null model constructed with 148 crAssphage genomes (they were downloaded using *ncbi-genome-download* from viral database with parameters: -s genbank, -l “all” --taxid 1978007). Network plots were generated using R packages *igraph*^78^ and *ggnetwork*. Genome diagram figures were prepared using the *GenoPlotR*^79^ package. Other plots were constructed using *ggplot2*^80^, *ggpubr*^81^ and *pheatmap*.

## Supporting information

Supplementary figures, tables, methods, and results

## Data availability

Sequencing data is available in the NCBI under BioProject PRJNA747701. Contigs of metagenome-assembled genomes (MAGs) and their CRISPR spacers are provided at DOI: 10.5281/zenodo.5122918.

## Code availability

All software used in this manuscript are currently published and available in the appropriate repositories. They are completely listed, as well as the setting used, in the Methods session.

## Acknowledgements

We thank the individual cheese producers and the Canastra Cheese Production Association (APROCAN) for their support to this research. We also thank Fabiana Lima for her invaluable lab benchwork work support. This work was supported by the Brazilian National Research Council (CNPq), grant: 425157/2018-0, from the Sao Paulo Research Foundation (FAPESP), grant: 2013/07914-8, and the Coordenação de Aperfeiçoamento de Pessoal de Nível Superior - Brasil (CAPES) - Finance Code 001.

## Author contributions

L.L.Q., G.A.L. and C.H. conceived of the project. L.L.Q. and W.R.I. performed experiments. L.L.Q. and C.H. analysed data and wrote the manuscript. G.A.L., M.L., U.M.P., B.D.G.M.F. and C.H. contributed funding. All authors reviewed and approved the manuscript.

