## Supplementary figures, tables, methods, and results for "High Level of Interaction between Phages and Bacteria in an Artisanal Raw Milk Cheese Microbial Community"

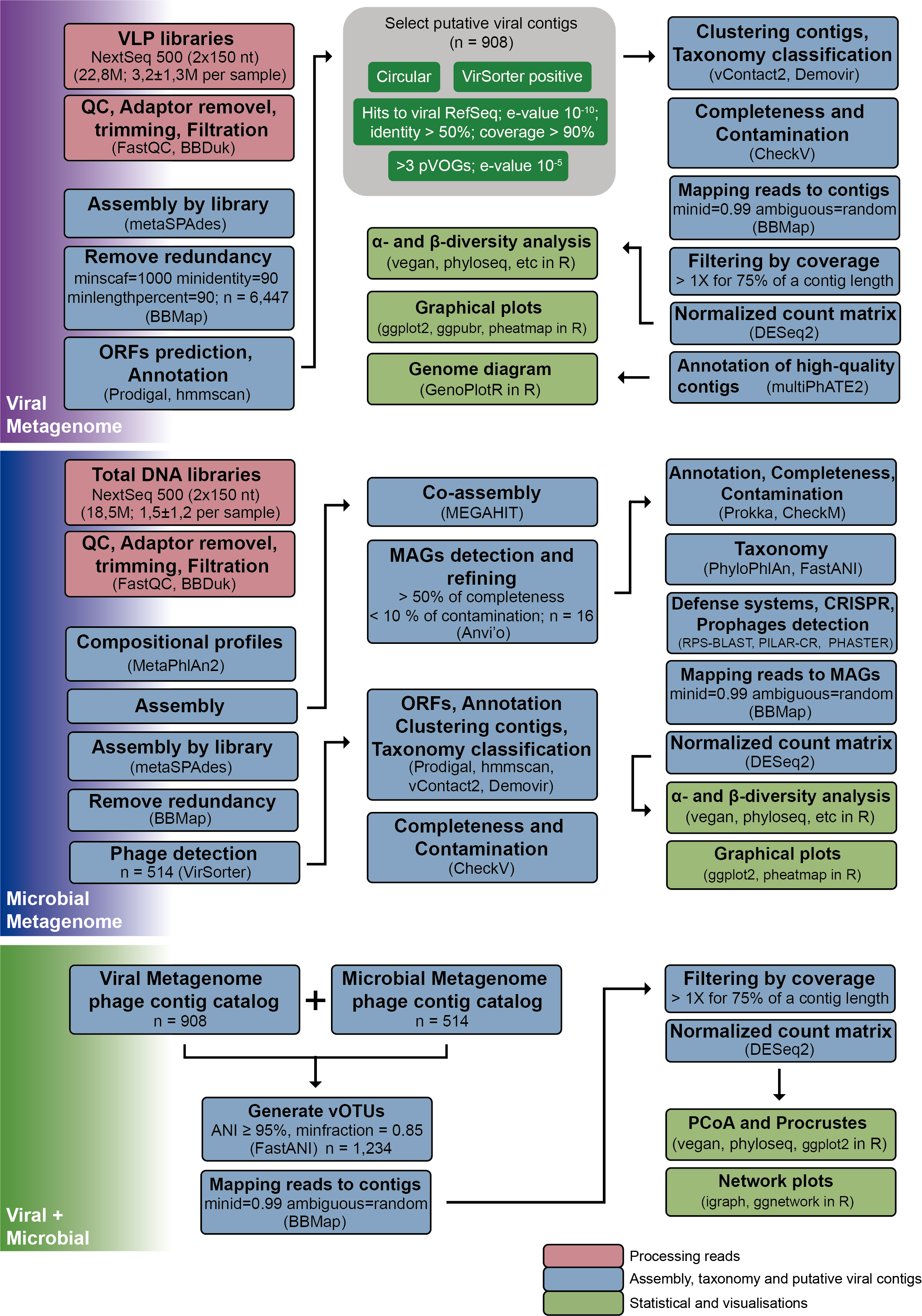

**Fig S1.** **Bioinformatics pipeline and statistics analysis used for processing viral metagenome (VLPs) and microbial metagenome.**

**Fig S2.** **Viral contigs recovered by viral metagenome (VLP) and classified as prophage by VirSorter.** Size and coverage distribution of 908 putative viral genomes detected (coverage depth by genome length in bp) and which were classified as prophage by VirSorter.

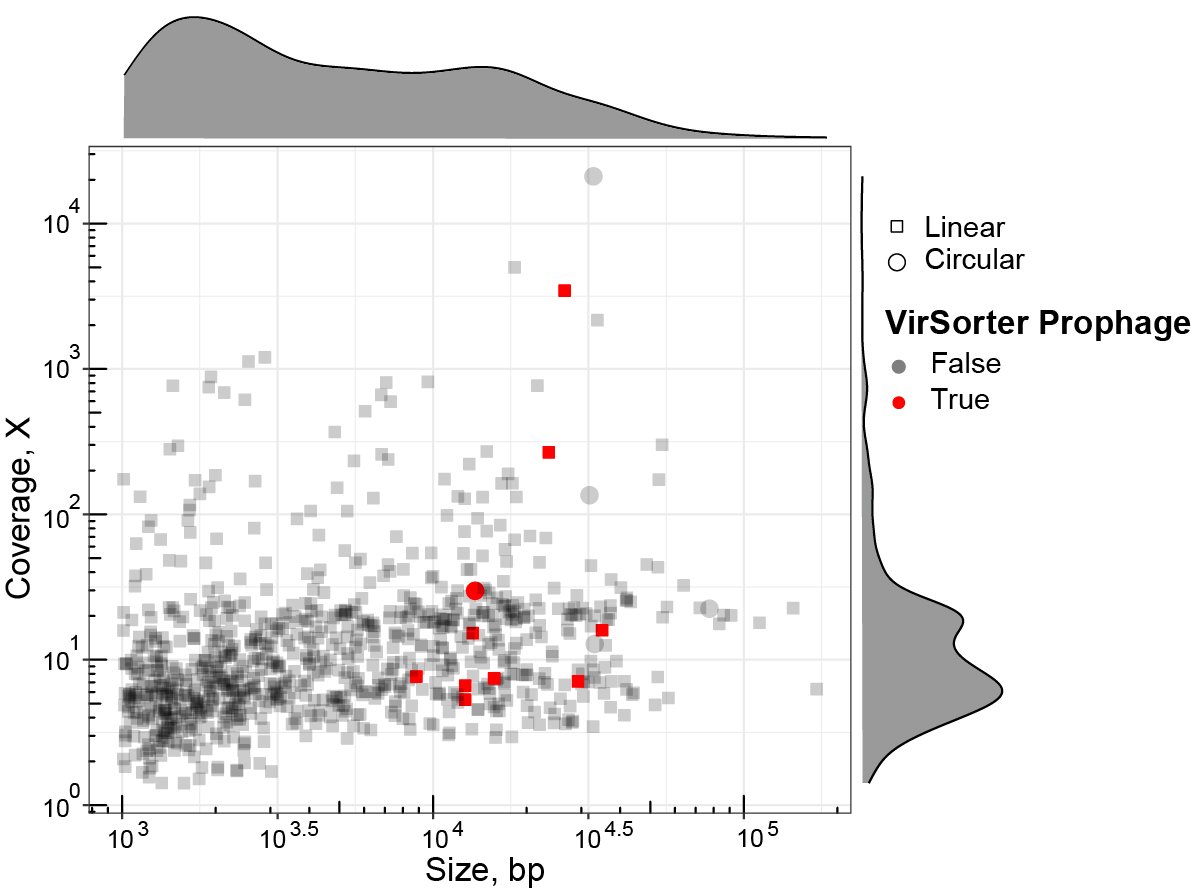

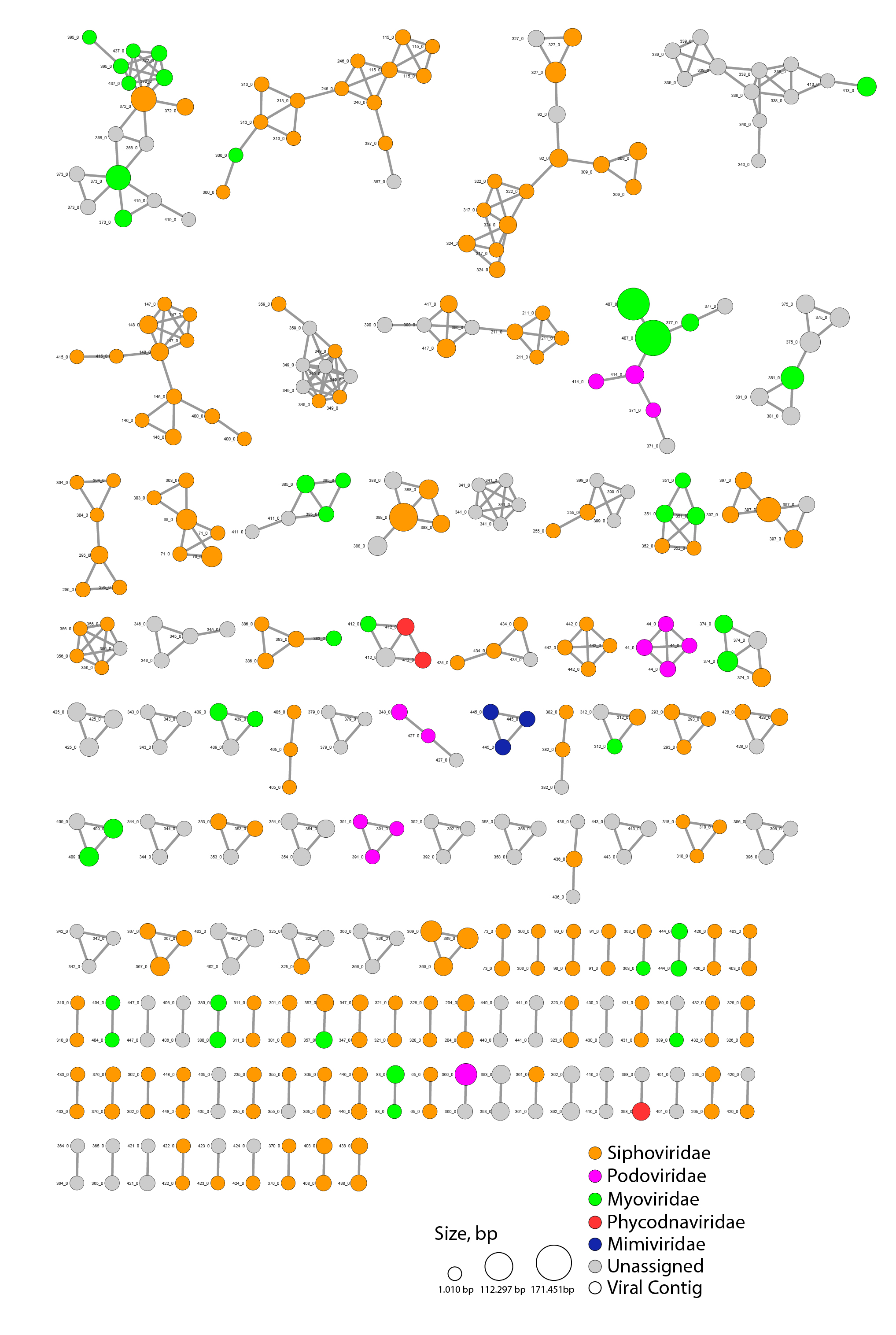

**Fig S3.** **Viral Clusters (VC) formed between viral contigs.** Viral contigs and their family level classification are represented by circles. Colors indicate viral families and the size of circles represent the viral contig length in base pairs.

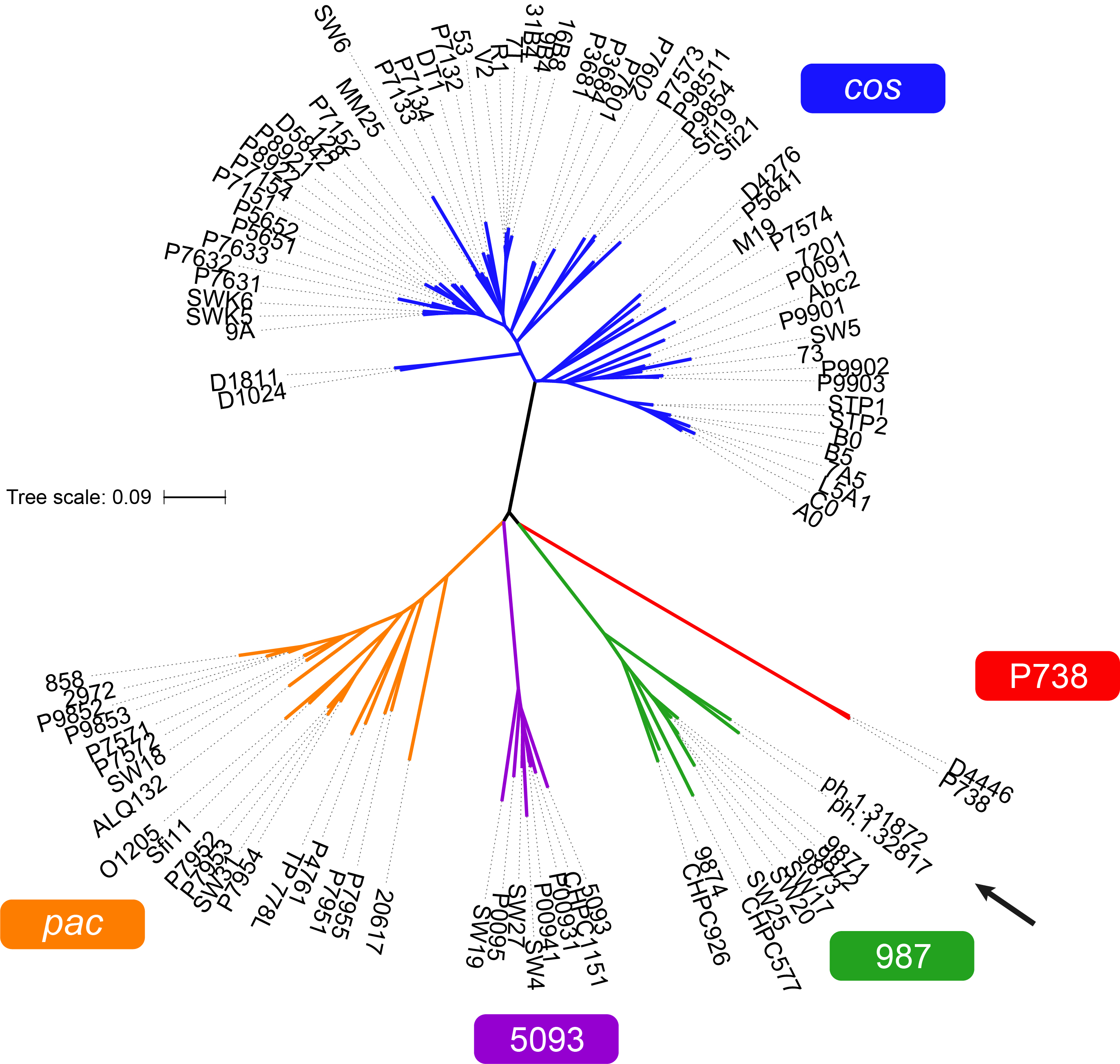

**Fig S4. Phylogenomic tree of *Streptococcus* phages.** Colors represent the genus of phages and the arrow indicates the genomes of phages ph.1.31872 and ph.1.32817 located within genus 987 group.

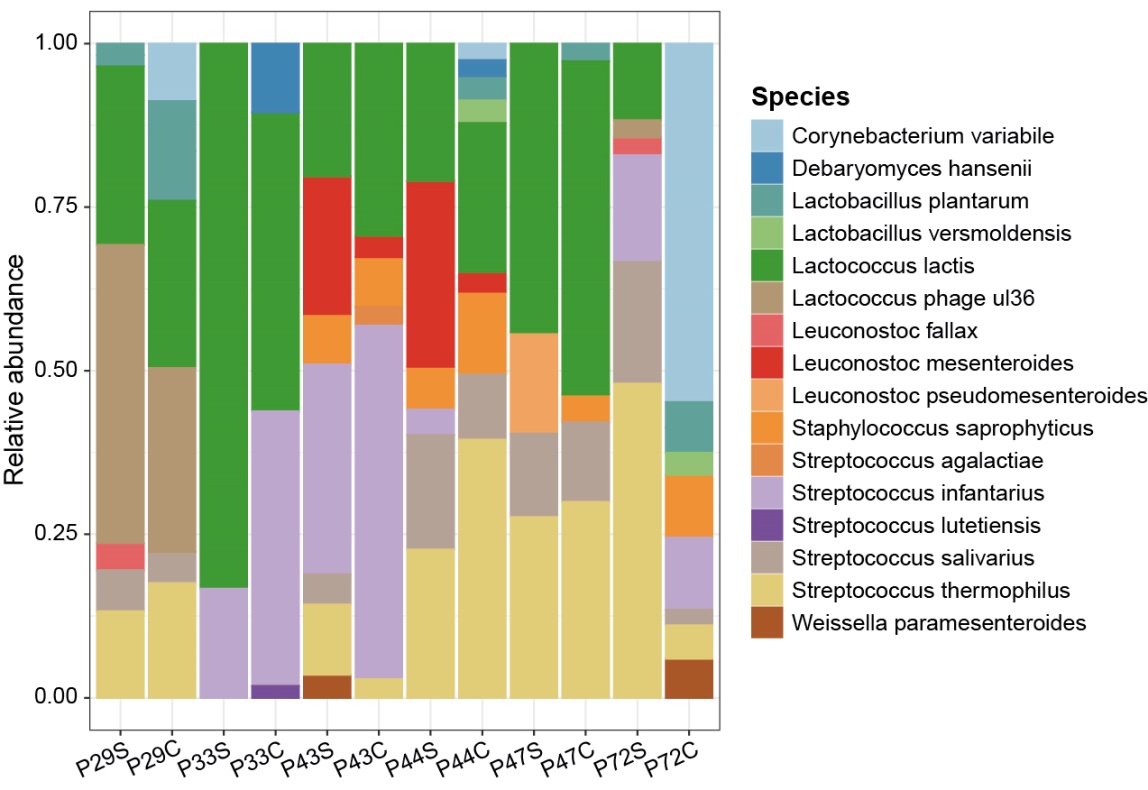

**Fig S5. Compositional profiles of microbial metagenomes in endogenous starter culture and cheese.** Relative abundance of each species in 12 paired starter culture (e.g. P29S) and cheese samples (e.g. P29C) assessed using MetaPhlAn2**.**

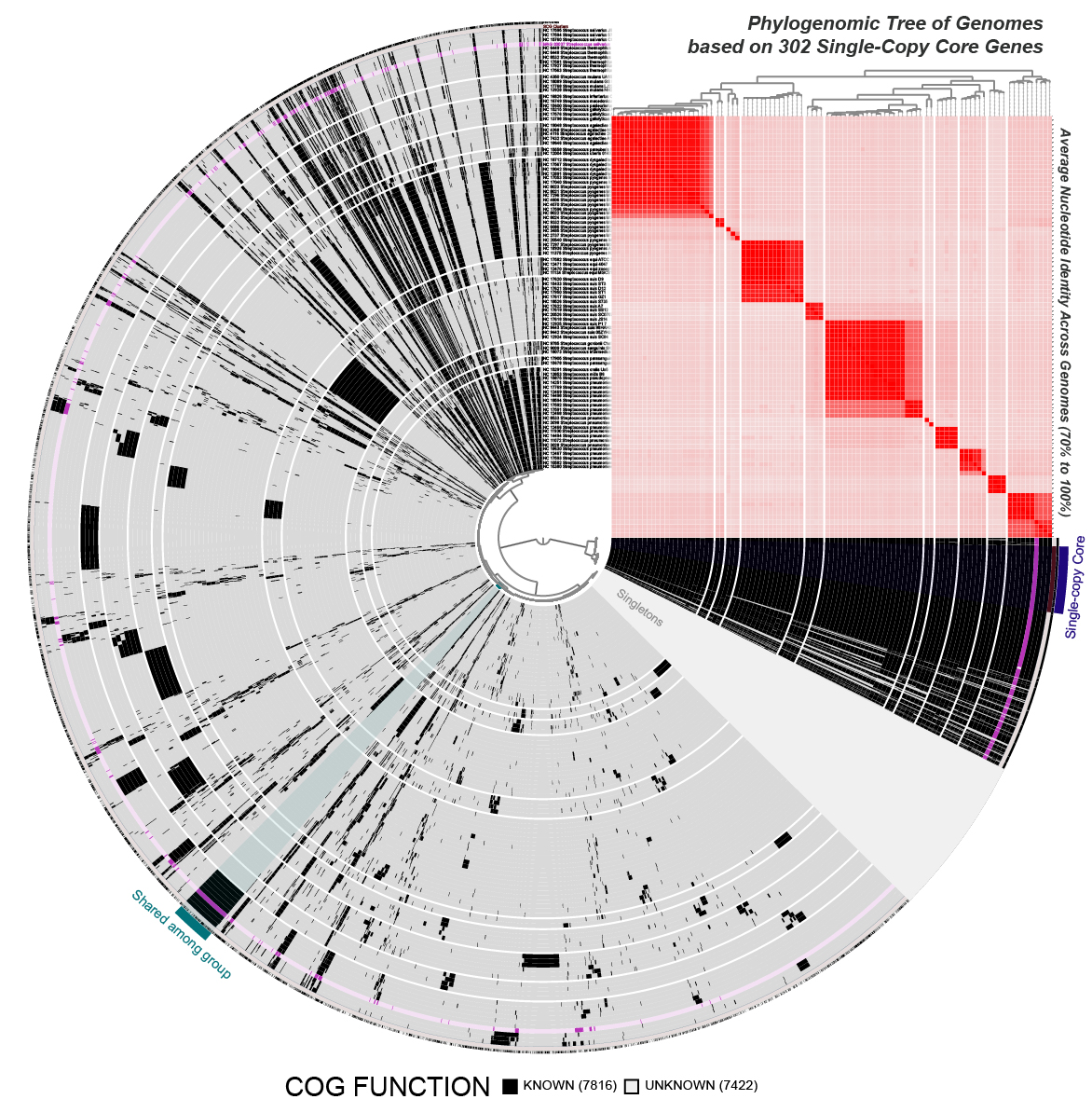

**Fig S6. Pangenomic analysis of *Streptococcus* genus.** It was performed with 94 complete genomes selected from previous study and metagenome-assembled genomes classified as *Streptococcus salivarius* (MAG7; highlighted in purple).

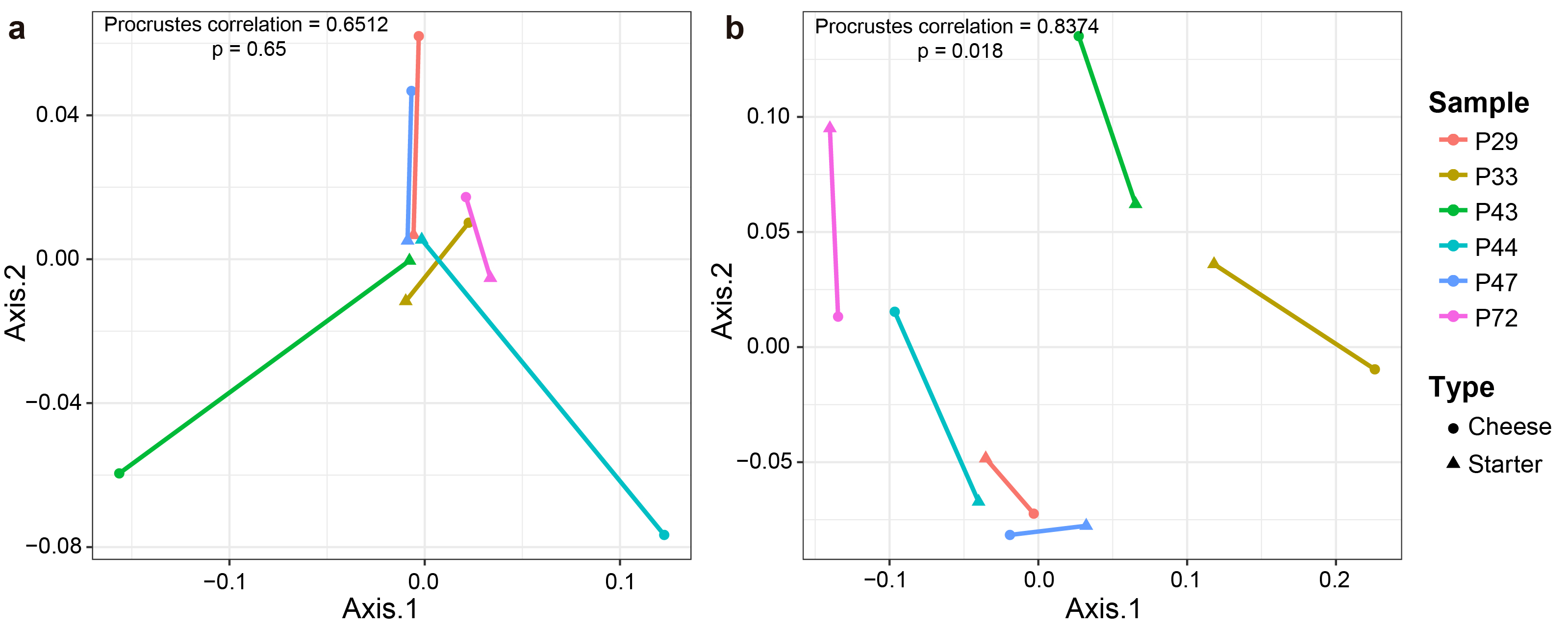

**Fig S7. Procrustes analysis using PCoA coordenations of starter culture and cheese communities.** **a**, Correlations between viral community in starter cultures and cheese samples. **b**, Correlations between bacterial communities in starter cultures and cheese samples. Distance matrices were calculated using Jensen-Shannon divergence in both cases.

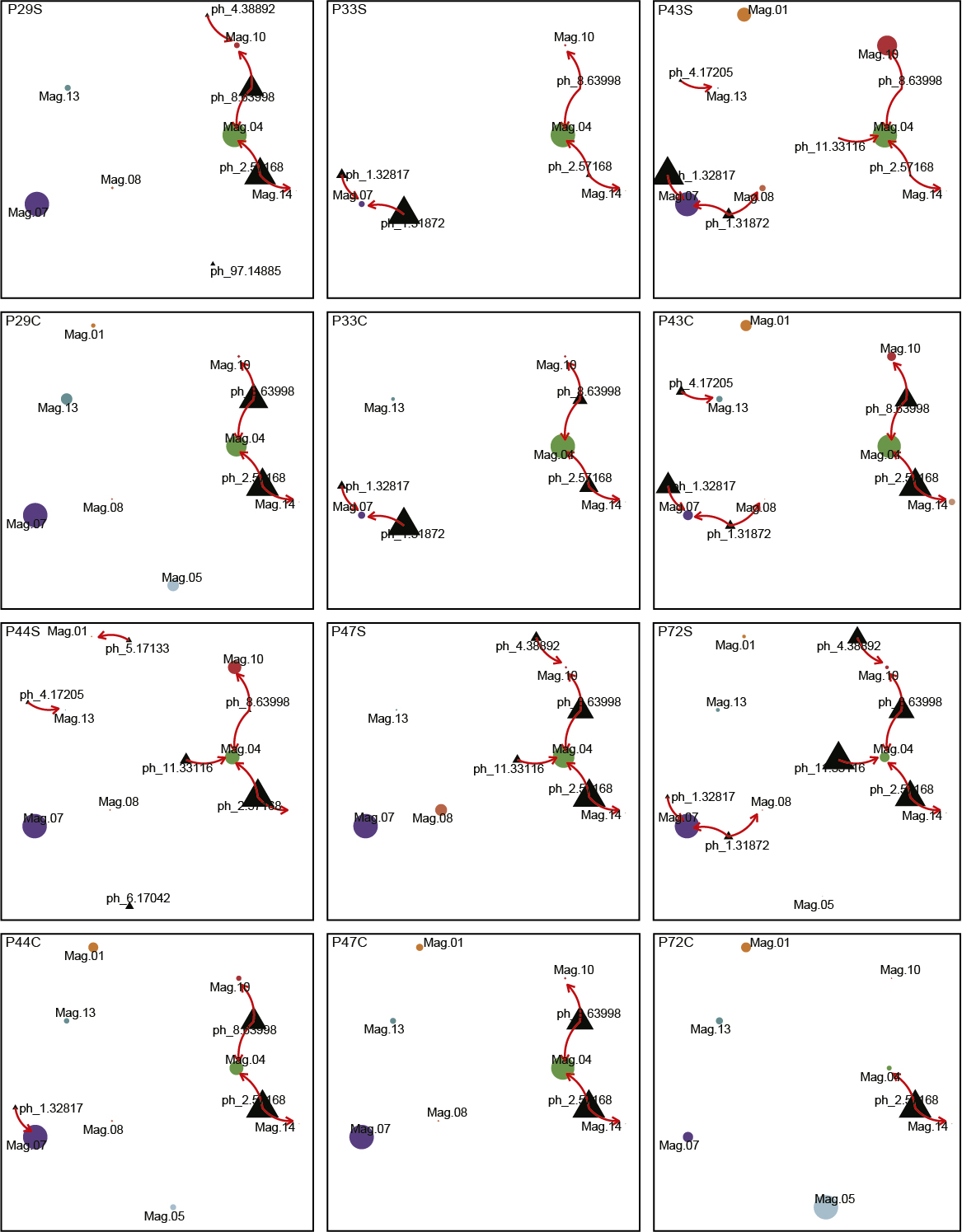

**Fig S8. Phage-bacteria interaction network during cheese production.** Putative phage-bacteria infection interactions for all producers (e.g. P29 and P33), and for separate starter and cheese samples (e.g. P29S and P29C). Nodes represent each MAG and phage analysed, edges represent putative infection relationships between phage and bacteria. This figure refers to Fig. 5.

**Supplementary Tables**

The supplementary tables with more than 10 columns or rows were submitted separately (Table S1, S3-S7, and S9-S11).

**Table S2**. Alpha-diversity of viral metagenome

| sample | reads | observed | shannon | simpson |
| --- | --- | --- | --- | --- |
| P29 | 3561923 | 759 | 6.83812 | 0.94896 |
| P33 | 53071 | 108 | 2.64524 | 0.53693 |
| P43 | 7500568 | 108 | 1.29953 | 0.39765 |
| P44 | 438574 | 465 | 6.73909 | 0.97449 |
| P47 | 1493347 | 384 | 4.66591 | 0.78968 |
| P72 | 37684 | 286 | 7.46798 | 0.992 |
| P85 | 234953 | 313 | 7.35305 | 0.98982 |

**Table S8.** Significant correlations between phage and MAG abundances from starter to cheese

| Phage | MAG | cor | pval |
| --- | --- | --- | --- |
| NODE_11_length_33116_cov_15.986812 | CANASTRA_MAG_00004 | -0.845154255 | 0.034109423 |
| NODE_1_length_31872_cov_135.581324 | CANASTRA_MAG_00007 | -0.880406274 | 0.020598735 |
| NODE_1_length_32817_cov_14173.771931 | CANASTRA_MAG_00007 | -0.880406274 | 0.020598735 |
| NODE_1_length_31872_cov_135.581324 | CANASTRA_MAG_00008 | -0.83165549 | 0.04012438 |
| NODE_8_length_63998_cov_19.026352 | CANASTRA_MAG_00010 | -0.885714286 | 0.018845481 |
| NODE_2_length_57168_cov_11.104757 | CANASTRA_MAG_00014 | -0.885714286 | 0.018845481 |
| NODE_1_length_32817_cov_14173.771931 | CANASTRA_MAG_00007 | -0.845154255 | 0.034109423 |
| NODE_4_length_38892_cov_19.980174 | CANASTRA_MAG_00010 | -0.880406274 | 0.020598735 |
| NODE_4_length_17205_cov_17.695569 | CANASTRA_MAG_00013 | -0.845154255 | 0.034109423 |

**Supplementary Methods**

List of phage-specific defense systems used to manual filtering identification of marker genes in MAGs genomes. This list was based on Bezuidt et al.^1^.

| **R.M** | *hsdR*, *hsdM/hsdS*, *yhdJ, ssl2, mcrA, yeeA , mrr, COG2810, mcrBC* |
| --- | --- |
| **DISARM** | *drmA, drmB, drmC , drmD*, *drmMI, drmMII* |
| **BREX** | *brxA, brxC*, *brxHI, brxL, brxP, pglX*, *pglW*, *pglZ* |
| **Druantia** | *druE,* *druM* |
| **Abi** | *abiEi*, *abiEii*, *abiF*, *abiTii*, *abiH*, *abiU2*, *abiJ*, *abiG*, *abiA*, *abiV*, *abiGi*, abiGii, abiC |
| **Zorya** | *zorA*/*zorB, zorC,* zorD, zorE |
| **Septu** | *ptuA*, *ptuB* |
| **Gabija** | *gajA,* *gajB* |
| **Theoris** | *thsA, thsB* |
| **CRISPR-cas Type I** | *cas3 , cas5, csp1, csp2* |
| **Type I-A** | *cas8a1, csx13* |
| **Type I-B** | *cas1-HMARI*, *cas1-MYXAN, csh2, cst2, cmx8* |
| **Type I-C** | *cas5d, csd1, csx17* |
| **Type I-D** | *csc1, csc2, csc3, cas10d, cas8c* |
| **Type I-E** | *cse1, cse2, cas5e, PRK13921* |
| **Type I-F** | *csy1, csy2, csy3, csy4, cas1-YPEST* |
| **Type I-U** | *GSU0052, GSU0053, GSU0054, csb1, csb2, csb3, csx15* |
| **CRISPR-cas Type II** | *cas1, csn1, cas-NMENI, cas9* |
| **Type II-B** | *csx12* |
| **CRISPR-cas Type III** | *cas10, cas6, csx1, csx3* |
| **Type III-A** | *csm1, csm2, csm3, csm4, csm5, csm6, TM1806* |
| **Type III-B** | *cmr1, cmr3, cmr4, cmr5, cmr6, csx1* |
| **Type III-BC** | *cmr1, cmr3, cmr4, cmr5, cmr6, csx1* |
| **Type III-D** | *csx10* |

**Supplementary Results**

**Bacteriophage taxonomy classification.** Using CheckV, as well as the Minimum Information about an Uncultivated Virus Genome (MIUViG) criterium^2^, we classified our genomes as: complete (5), high-quality (12), medium-quality (23), low-quality (584), and not-determined quality (284) (Table S1). We further refined the classification of our viral catalog by comparing it to the RefSeq complete viral genome database using BLAST and stringent criteria (e-value < 10^-10^, coverage > 90% of contig length and > 50% identity), and obtained 94 viruses classified at species level. The detected viruses were *Lactococcus* phages 949, asccphi28, bIL285, bIL286, bIL309, bIL312, P078, P162 and ul36. We also found *Lactobacillus* phages, such as phiAQ113 and phiJL1; *Staphylococcus* phages GRCS and phiSA12; and *Streptococcus* phages 9872 and 9874 (for a detailed list, please see Table S1).

Viral contigs clustered with RefSeq genomes belong to *Siphoviridae*, *Myoviridae* and *Podoviridae* families and their sizes ranged from 1,001 to 54,569 bp. We observed that some VCs were composed of complete (VC 148: *Streptococcus virus*) and high-quality (VC 248: *Staphylococcus phage*; VC 44: *Actinomyces virus*; VC 67: *Arthrobacter*, *Gordonia*, and *Rhodococcus phages*; and VC 69: *Lactococcus phage*) viral contigs. Also, a high-quality genome clustered with 7 RefSeq genomes of *Staphylococcus phages* (i.e. GRCS) that belong to the genus *Rosenblumvirus* (VC 248).

The ANI measured between ph.72.18300 (VC320) and asccphi28 genomes was 93%, well below the usually accepted threshold for same species classification, indicating that this phage is potentially a novel viral species within the *Lactococcus* phage asccphi28 group. This phage was detected in high abundance in only one of the studied producers.

**Profage characterization.** Four intact prophage sequences were identified in MAGs classified as *Leuconostoc fallax* (MAG2), *Escherichia coli* (MAG3), *Streptococcus infantarius* (MAG6), and *Streptococcus salivarius* (MAG7). All these sequences were classified at family level as *Siphoviridae*. The most common gene annotations produced by PHASTER were *Lactobacillus* phage Sha1 (7 genes) for the prophage found in MAG2, *Salmonella* phage SEN34 (6) for the prophage in MAG3, *Streptococcus* phage phiNJ2 (20) for the prophage in MAG6, and *Streptococcus* phage 5093 (9) for the prophage in MAG7 (Table S7).
